# Mechanism of the Dual Action Self-Potentiating Antitubercular Drug Morphazinamide

**DOI:** 10.1101/2024.10.08.617272

**Authors:** Lev Ostrer, Taylor A. Crooks, Michael D. Howe, Sang Vo, Ziyi Jia, Pooja Hegde, Nathan Schacht, Courtney C. Aldrich, Anthony D. Baughn

**Author notes:** Correspondence can be made to Anthony D. Baughn. **Email:**. These authors contributed equally to this work (Lev Ostrer, Taylor A. Crooks).

## Abstract

Pyrazinamide (PZA) is a cornerstone of first-line antitubercular drug therapy and is unique in its ability to kill nongrowing populations of *Mycobacterium tuberculosis* through disruption of coenzyme A synthesis. Unlike other drugs, PZA action is conditional and requires potentiation by host-relevant environmental stressors, such as low pH and nutrient limitation. Despite its pivotal role in tuberculosis therapy, the durability of this crucial drug is challenged by the emergent spread of drug-resistance. To advance drug discovery efforts, we characterized the activity of a more potent PZA analog, morphazinamide (MZA). Here, we demonstrate that like PZA, MZA acts in part through impairment of coenzyme A synthesis. Unexpectedly, we find that, in contrast to PZA, MZA does not require potentiation and maintains bactericidal activity against PZA-resistant strains due to an additional mechanism involving aldehyde release. Further, we find that the principal mechanism for resistance to the aldehyde component is through promoter mutations that increase expression of the mycothiol oxidoreductase MscR. Our findings reveal a dual action synergistic mechanism of MZA that results in a faster kill rate and a higher barrier to resistance. These observations provide new insights for discovery of improved therapeutic approaches for addressing the growing problem of drug-resistant tuberculosis.

**Significance Statement:** Pyrazinamide is the only antitubercular drug of its kind, capable of targeting persistent *Mycobacterium tuberculosis* through disruption of the coenzyme A biosynthetic pathway. With the emergent spread of drug-resistant tuberculosis, it is imperative to identify more effective next-generation drugs. In this study, we characterized the mechanism of action of a more potent analog of pyrazinamide, morphazinamide. We demonstrate that like pyrazinamide, morphazinamide impairs coenzyme A metabolism. In contrast to pyrazinamide, we find that morphazinamide has an additional aldehyde-dependent mechanism that mediates potent bactericidal activity against both pyrazinamide-susceptible and pyrazinamide-resistant strains of *M. tuberculosis*. These findings open up new opportunities for the development of next-generation antitubercular drugs to tackle the increasing challenge of drug-resistant tuberculosis.

## Introduction

For several decades, pyrazinamide (PZA; Figure 1A) has been an integral part of the first-line standard short course therapy for tuberculosis (TB) (1). A unique feature of PZA is its ability to kill non-replicating populations of the causative agent *Mycobacterium tuberculosis* (2).

**Figure 1.**
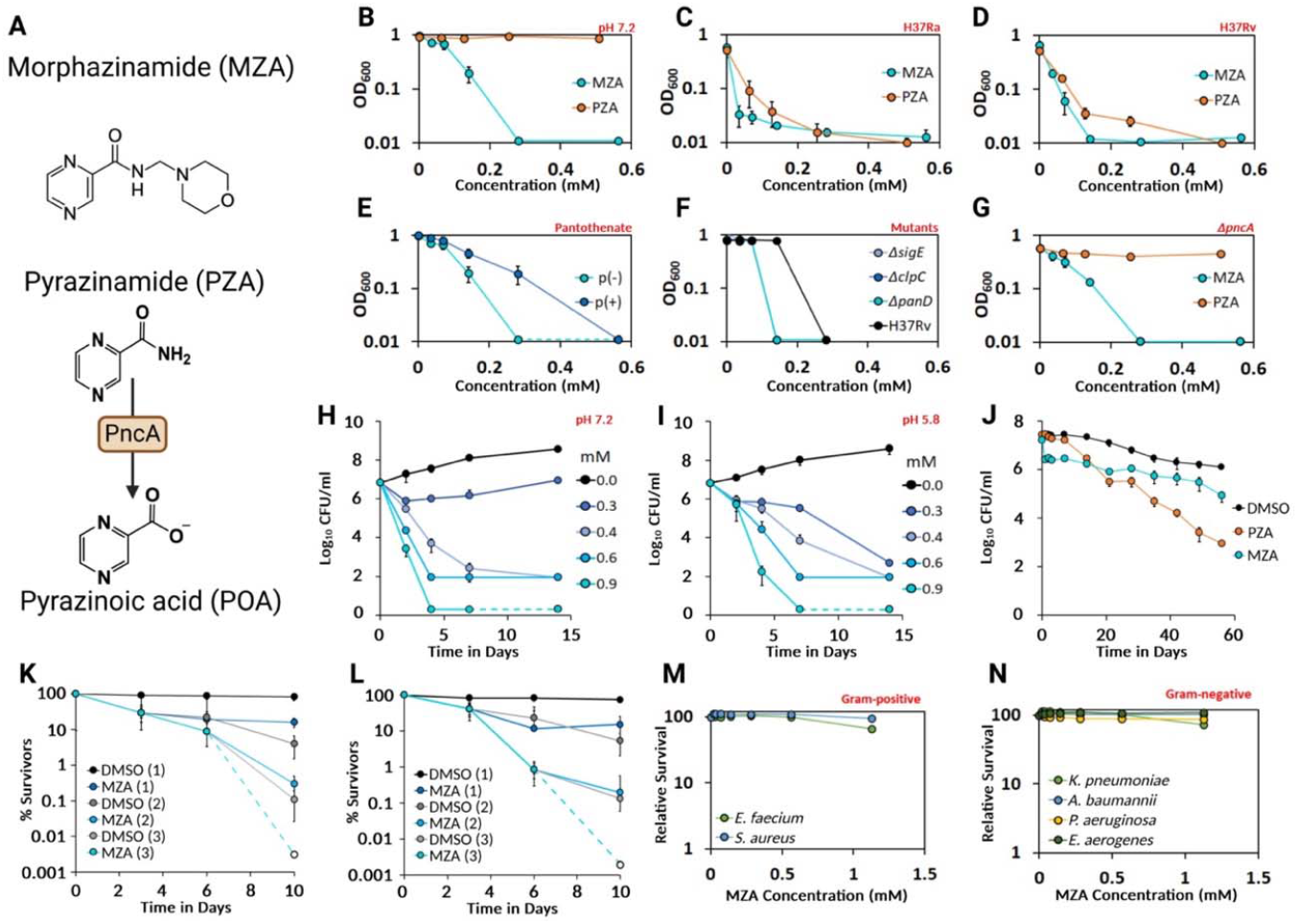
Impact of MZA on PZA susceptible and resistant *M. tuberculosis*. **A**, Structure of MZA and activation of PZA to POA by PncA_Mtb_. Dose response curves comparing MZA and PZA susceptibility for *M. tuberculosis* strains **B**, H37Ra at pH 7.2 and **C**, H37Ra at pH5.8 and **D**, H37Rv at pH 5.8. **E**, Assessment of pantothenate antagonism of MZA action on H37Ra at pH 7.2. Determination of MZA susceptibility of PZA resistant strains **F**, H37Rv Δ*sigE*, H37Rv P_*clpC1*_::*himar1*, H37Rv *panD*::*himar1* and **G**, H37Rv Δ*pncA* compared to parental H37Rv. Dose dependent kill curves of H37Rv exposed to MZA at **H**, pH of 7.2 and **I**, pH 5.8 over a 14-day period. **J**, Kill curves performed with H37Rv under starvation conditions following a one-time treatment with PZA, MZA or DMSO over a 60-day period at pH of 7.2. Kill curves performed with **K**, H37Rv and **L**, H37Rv Δ*pncA* (PZA resistant) incubated at neutral pH under starvation conditions with MZA replenished every 3 days. Evaluation of MZA activity against **M**, Gram-positive and **N**, Gram-negative ESKAPE pathogens. All assays were performed in biological triplicate with mean displayed and error-bars indicative of standard deviation.

Use of PZA has played a pivotal role in reducing TB relapse rates and shortening the duration of treatment from 9 to 6 months (1). In spite of this success, wide use of this treatment regimen has resulted in the emergent spread of PZA resistance. Current rates of PZA resistance in *M. tuberculosis* clinical isolates range from 16 to 42% depending upon patient cohort (3). The primary mechanism of clinical PZA resistance occurs through spontaneous loss-of-function mutations in the *M. tuberculosis pncA* gene which encodes an amidase that is required for activation of PZA to pyrazinoic acid (POA; Figure 1A) (4, 5). Identification of potent next-generation analogs of PZA that circumvent resistance will be critical to maintain use of this important therapeutic tool.

Despite its crucial role in TB therapy the antitubercular mechanism of PZA action is not fully defined. Recent evidence demonstrates that PZA acts through disruption of synthesis of the thiol cofactor coenzyme A (CoA), in part, through destabilization of L-aspartate decarboxylase (PanD) by POA (6, 7). Intriguingly, while CoA plays an indispensable role in *M. tuberculosis* metabolism, PZA susceptibility is conditional and requires activation of the cell envelope stress response through exposure of bacilli to low pH or a variety of other stressors (8). How disruption of CoA biosynthesis connects with conditional susceptibility of *M. tuberculosis* to PZA has not been resolved (9). It has recently been shown that exposure of *M. tuberculosis* to low pH results in thiol stress (10), yet, whether thiol stress is associated with the enhanced impact of PZA on CoA metabolism has not been reported. Consistent with an essential role for environmentally driven potentiation of PZA action *in vivo*, PZA shows no detectable antitubercular activity in mice that lack T cells and are thereby impaired for macrophage activation which is required for full phagosomal acidification (11). Identification of means to bypass the need for host-mediated potentiation of PZA action would advance antitubercular therapy through promoting drug potency in the context of compromised immunity which is causally associated with progression of TB (9).

To address the unmet needs outlined above, we set out to characterize the mechanism of action of an intriguing PZA analog, morphazinamide (MZA, Figure 1A). Early clinical studies demonstrated efficacy of MZA against *M. tuberculosis* in humans with equivalent potency and safety profiles relative to PZA (12, 13). Further, *in vitro* assays indicated that the antitubercular activities of MZA and PZA were comparable, with similar levels of growth inhibition in broth culture (14). However, it was noted that MZA and other aminomethylene analogs did not require low pH to show inhibitory activity and retained activity against PZA resistant isolates, suggesting that they operate by a distinct mechanism (14). In this study we applied approaches in chemical biology, functional genomics and bacterial physiology to characterize the mechanism of action of MZA. Similar to its sustained activity against *pncA* null strains of *M. tuberculosis*, we find that MZA also retains activity against *M. tuberculosis* strains with newly described PZA resistance mechanisms (8). By using transcriptional profiling and transposon sequencing (Tn-seq) we find many parallels between responses of *M. tuberculosis* to PZA, POA and MZA that highlight the impact of these drugs on CoA metabolism. Importantly, we uncover an additional mechanism unique to MZA involving aldehyde release and find that MZA susceptibility in particular is strongly influenced by the mycothiol oxidoreductase MscR. Together, our findings illuminate new opportunities to improve potency and raise the resistance barrier to PZA analogs.

## Results

### Impact of MZA on PZA susceptible and resistant *M. tuberculosis*

To assess the antitubercular activity of MZA, we performed minimum inhibitory concentration assays (MICs) using a panel of strains with differing mechanisms of PZA resistance and under various conditions that influence PZA susceptibility. We began by comparing activity between PZA and MZA using standard growth conditions at circumneutral pH with both H37Ra and H37Rv backgrounds. Consistent with the known conditional susceptibility of *M. tuberculosis* to PZA, cultures grew unimpaired at concentrations as high as 4.5 mM in medium at circumneutral pH (Figure 1B; Table S1). In striking contrast, under these same conditions we observed >90% inhibition of growth in the presence of 280 µM MZA (Figure 1B; Table S1). Next, we assessed whether MZA activity could be enhanced by conditions that potentiate PZA and diminished under conditions that antagonize PZA. To evaluate potentiation, MICs were determined under acidic (pH 5.8) conditions (Figure 1C,D). To determine if MZA activity could be antagonized similarly to PZA, the effect of the CoA intermediate pantothenate on drug action was evaluated. Consistent with previous work, PZA was fully inhibitory at 250 µM under acidic conditions (Figure 1C,D; Table S1), while the addition of 230 µM pantothenate fully antagonized PZA action (15) (Figure 1E; Table S1). Acidic conditions were able to enhance MZA activity by four-fold, lowering its MIC to 70 µM (Figure 1C,D; Table S1), whereas addition of pantothenate caused a two-fold antagonistic effect on MZA activity at both acidic and circumneutral pH (Figure 1E; Table S1). These observations indicate that MZA is mildly impacted by conditions that strongly influence PZA action, suggesting key mechanistic differences despite underlying molecular similarity between these two drugs.

We next sought to determine whether MZA retained activity against previously described PZA resistant isolates. These strains included mutants deleted for *pncA* (required for activation of PZA to POA) and *sigE* (required for potentiation of PZA susceptibility at low pH), as well as a strain with truncation of the 3’ end of *panD* (8), which is thought to interfere with POA action through stabilization of PanD (7). Despite PZA showing diminished activity against these strains, MZA maintained activity with MICs ranging from 140 to 280 µM (Figure 1F,G; Table S1). It is worth noting that the Δ*pncA* strain showed a two-fold increase in its MZA MIC relative to the parental strain (Figure 1G; Table S1) indicating a contribution of PncA to MZA action.

One of the key features of PZA is its unique ability to elicit a bactericidal effect on non-growing *M. tuberculosis* (16). Thus, we sought to determine whether MZA also has bactericidal activity against actively growing and metabolically quiescent drug tolerant bacteria by performing kill curves under nutrient rich and nutrient limiting conditions. When mid-exponential cultures under either acidic or circumneutral pH growth conditions were treated with 900 µM MZA, greater than 99.999% killing was observed after 4 days (Figure 1H,I). After 14 days of incubation following a single treatment, no surviving CFU were observed. We next tested the bactericidal activity of PZA and MZA under nutrient limiting conditions by exposing starved bacteria to either 900 µM PZA or MZA. After 56 days of stasis, the majority of untreated bacilli survived, whereas there was more than 99.9% loss in viability of bacilli exposed to 900 µM PZA (Figure 1J). In contrast to PZA, MZA bactericidal activity was transient, with 80% killing in the first 2 days followed by a plateau (Figure 1J). To determine whether the plateau in killing was due to drug inactivation or due to enrichment of a drug tolerant population, starved cultures were washed and treated with fresh MZA every 3 days. The number of survivors was normalized to the post-wash/pretreatment starting population after every round of treatment to account for any cells lost during the wash. As before, we observed 80% killing after 3 days for both H37Rv (Figure 1K) and H37Rv Δ*pncA* (Figure 1L). Following the second treatment over 99.5% of bacteria were no longer viable for both strains (Figure 1K,L). The third treatment resulted in sterilization of bacteria with no detectible CFU for either strain (Figure 1K,L). These data indicate that MZA has a potent PncA-independent bactericidal activity against both actively growing and metabolically quiescent *M. tuberculosis*.

To assess the antibacterial spectrum of MZA action, we evaluated its inhibitory activity against a panel of human-associated ESKAPE pathogens (17). All ESKAPE pathogens that were tested showed MICs greater than 10-fold higher than those for *M. tuberculosis* (Figure 1M,N), suggesting that like PZA, MZA activity is highly selective for mycobacterial species.

### Transcriptional responses of *M. tuberculosis* to PZA, POA and MZA exposure

To gain a better understanding of the impact of PZA, POA and MZA on *M. tuberculosis* physiology and to determine similarities and differences in global responses of bacilli to these drugs, we performed genome-wide transcriptional profiling using RNA-seq. Since PZA is only active at low pH, all treatments were performed using *M. tuberculosis* grown in acidic (pH 5.8) media. Cultures of exponentially growing H37Rv were exposed to 200 µM PZA, POA, MZA or vehicle only (DMSO) for 24 hours. The resulting transcriptional profiles revealed strikingly similar results (Figure 2A-C; Figure S1A). Specifically, we observed a significant overexpression of early genes in the CoA biosynthesis pathway, including *panB, panG, panC, panD and coaX* (Figure 2A-C). This observation is consistent with the current understanding that POA interferes with CoA biosynthesis through disruption of PanD activity (18, 19). Expression of most other genes of the CoA pathway were not greatly altered by exposure to PZA, POA and MZA with the exception of the final gene of the pathway *coaE* which was down-regulated by 1.5 to 2-fold relative to the no drug control. Due to the central role of CoA in lipid metabolism, we also observed differential expression of numerous genes involved in fatty acid synthesis (FAS) and utilization. The majority of genes associated with the FAS-I and FAS-II systems that are critical for long-chain and mycolic acid biosynthesis, respectively, were overexpressed under all three treatment conditions (Figure 2A-C,F). Interestingly, we also observed down regulation of the large gene cluster for synthesis and export of phthiocerol dimycocerosate (PDIM) virulence lipids, as well as differential expression of several other less well characterized genes potentially involved in polyketide and fatty acid synthesis (Figure 2F,G). Beyond lipid synthesis and metabolism, there were also indications of altered respiratory activity and oxidative stress with induction of genes for fumarate reductase, the microaerophilic-type cytochrome *bd* oxidase and catalase peroxidase *katG* (Figure 2F).

**Figure 2.**
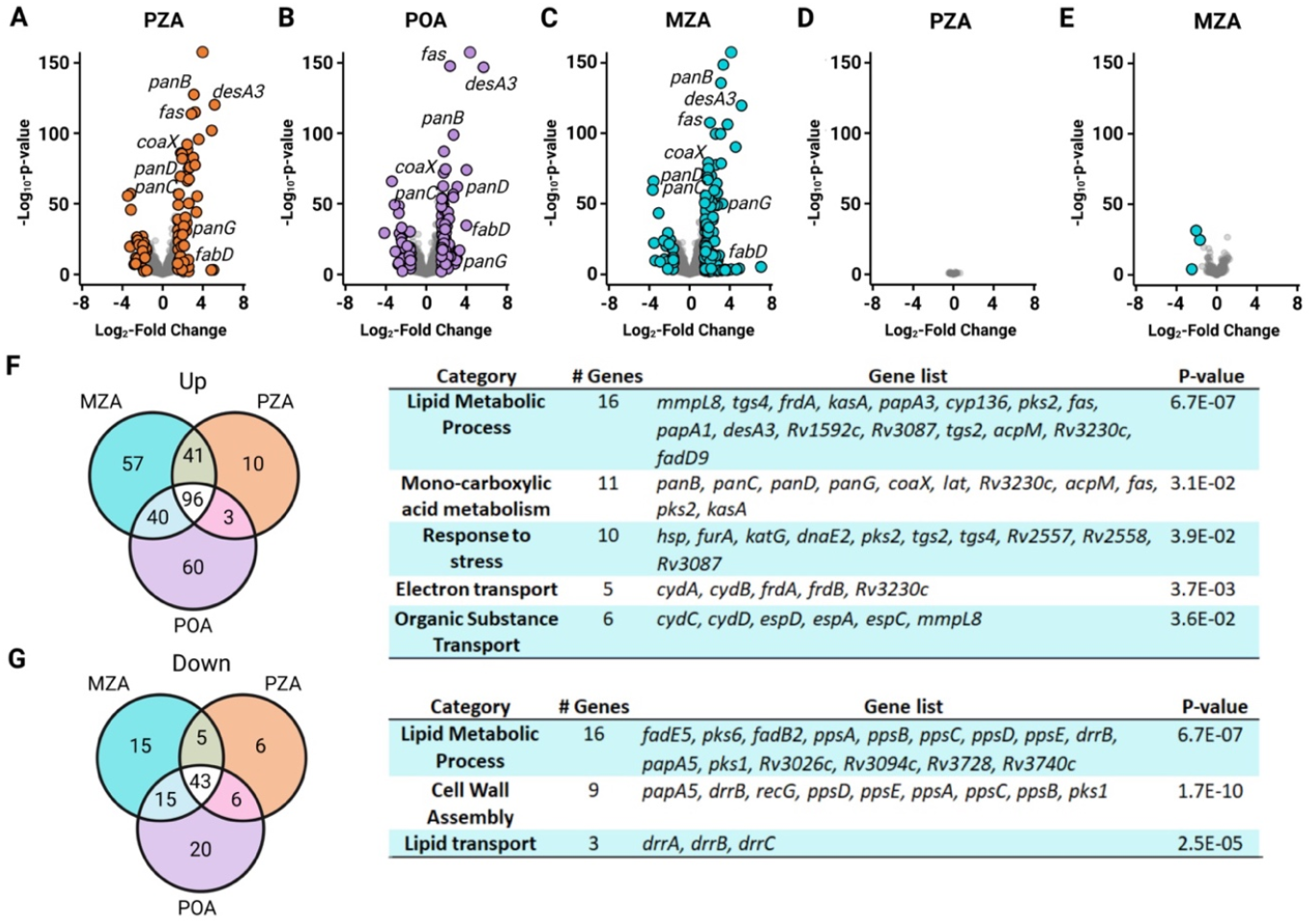
Transcriptional profiling of *M. tuberculosis* exposed to MZA, PZA and POA. Volcano plots showing significantly differentially expressed genes in presence of **A**, PZA, **B**, POA and **C**, MZA. Cells were treated with 200 µM MZA, POA or PZA at pH 5.8 for 24h prior to RNA purification and sequencing. The MZA, POA and PZA transcriptional profiles share many key features, including upregulation in coenzyme A and lipid metabolism pathways. Transcriptional changes associated with exposure of a H37Rv Δ*pncA* to **D**, PZA and **E**, MZA. Cells were treated with 200 µM MZA or PZA at pH 5.8 for 24h prior to RNA purification and sequencing. **F**, Venn diagram showing upregulated genes in MZA, POA and PZA treated cultures, with the corresponding GO term analysis of common genes to the right. **G**, Venn diagram showing downregulated genes in MZA, POA and PZA treated cultures, with the corresponding GO term analysis of common genes to the right. RNA-seq was performed in biological triplicate.

These similarities in responses of *M. tuberculosis* to MZA, PZA and POA exposure suggest some mechanistic conservation in their antimicrobial action. Our observation that deletion of *pncA* conferred a slight increase in the MIC for MZA led us to speculate that POA may be released from MZA in a PncA-dependent manner. To understand the role of PncA in response of *M. tuberculosis* to MZA, we performed RNA-seq using the H37Rv Δ*pncA* strain exposed to PZA or MZA (Figure 2D,E). As anticipated, the transcriptional profile of H37Rv Δ*pncA* showed no significant differences between the PZA-treated and untreated control (Figure 2D; Figure S1B). In contrast, the transcriptional profile of H37Rv Δ*pncA* exposed to MZA was distinct from the untreated control, suggesting that MZA exerts additional stress onto bacilli beyond PZA release (Figure 2E; Figure S1B). While the MZA response of H37Rv Δ*pncA* was relatively subdued compared to that of H37Rv, many of the differentially abundant transcripts included genes involved in amino acid synthesis, tRNA processing, ribosome assembly and protein chaperones, consistent with a response to proteotoxic stress.

### Release of PZA and POA from MZA

Based on the distinctions in antitubercular activity of PZA and MZA, and on the profoundly different transcriptional responses of H37Rv and H37Rv Δ*pncA* to MZA exposure, we hypothesized that MZA likely acts through two interacting mechanisms, one that is PncA-dependent and another that is PncA-independent. To parse these mechanisms, we assessed MZA metabolism in strains with and without PncA activity. We began by characterizing the rate of PZA release and conversion to POA. Exponentially growing H37Ra (PncA-proficient) cultures were amended with 250 µM MZA and abundance of MZA, PZA and POA was measured in cell extracts and media over an 8-hour time course via LC-MS. In the first 4 hours, cell-associated MZA rapidly accumulated then steadily declined, while PZA levels rose rapidly from 17 µM to 138 µM (Figure 3A). With continued incubation, levels of MZA and PZA declined to 34 µM and 121 µM, respectively. Meanwhile, limited POA accumulation was observed in cell lysates during the time course. The concentration of MZA present in the supernatant steadily declined from 250 to 100 µM over the eight-hour exposure, during which we also observed a steady increase of PZA from 4 to 67 µM and POA from 1 to 100 µM, respectively (Figure 3B).

**Figure 3.**
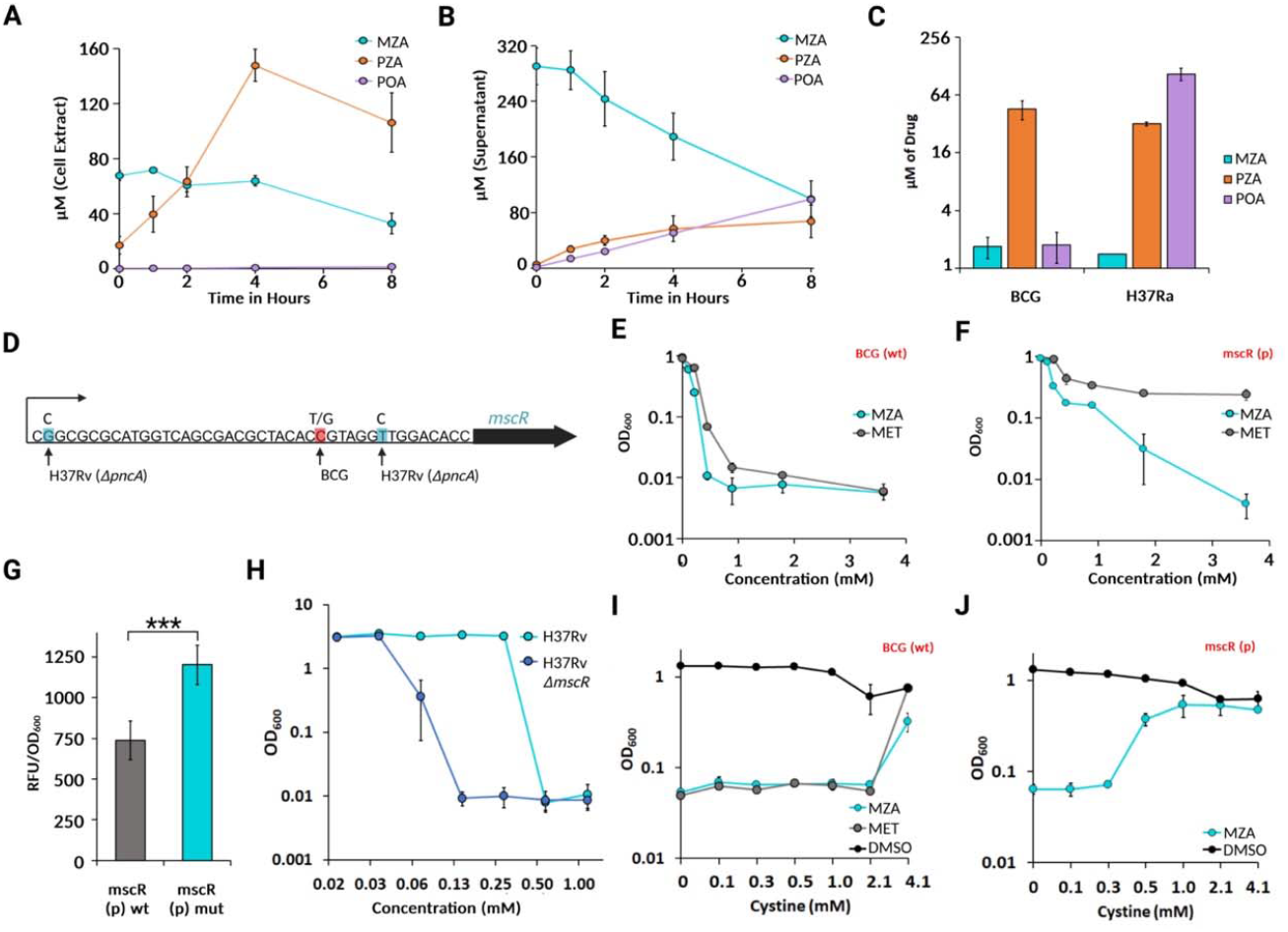
Metabolism of MZA results in release of PZA and drives thiol stress. Time course monitoring **A**, intracellular and **B**, extracellular abundance of MZA to PZA and POA from H37Ra exposed to 250 µM MZA. **C**, Intracellular abundance of MZA, PZA and POA following a 24-hr treatment of H37Ra (PZA susceptible) and *M. bovis* BCG (PZA resistant) with MZA. **D**, *mscR* promoter region highlighting MZA resistance mutations in H37Rv Δ*pncA* (blue) and BCG (red) backgrounds, respectively. Inhibition curve of an **E**, wild-type BCG and a **F**, *mscR* promoter mutant exposed to MZA and methenamine (MET). **G**, Comparison of GFP expression between wild-type *mscR* promoter and *mscR* promoter variant GFP reporter constructs in BCG. All samples were normalized to the OD_600_, significance was determined using a 2-tailed Student’s t-test using *n*=6. **H**, Inhibition curve comparing susceptibility of H37Rv and H37Rv Δ*mscR* to MZA. **I**, BCG wild-type and **J**, BCG *mscR* promoter mutant exposed to 0.9 mM MZA or MET in the absence or presence of cystine. All assays were performed in biological triplicate unless indicated otherwise with mean displayed and error-bars indicative of standard deviation.

To determine whether PncA is essential for formation of POA from MZA, a similar experiment was performed with overnight exposure using the vaccine strain *M. bovis* BCG which lacks pyrazinamidase activity due to a loss-of-function mutation in *pncA*. Following overnight exposure, cell-associated MZA was near the limit of detection for both BCG and the control strain H37Ra. Both strains showed accumulation of high levels of PZA, whereas only H37Ra showed an abundant level of POA (Figure 3C). Together, these data demonstrate a rapid conversion of MZA to PZA, followed by PncA-dependent conversion of PZA to POA.

### MZA impacts thiol metabolism

To further characterize the PncA-independent mechanism of MZA action, we selected for spontaneously resistant mutants of H37Rv, H37Rv Δ*pncA* and BCG. Selections were performed using solid medium containing a geometric series of concentrations from 280 to 900 µM MZA. No resistant isolates were recovered from the H37Rv background at any of the tested concentrations, nor for strains H37Rv Δ*pncA* and BCG in the presence of 900 µM MZA. In contrast, H37Rv Δ*pncA* and BCG strains yielded mutants resistant to 410 µM MZA at a frequency of 10^-5.9^, and to 610 µM MZA at a frequency of 10^-7^. Since *pncA* inactivation is necessary for isolation of MZA resistant isolates, the PZA-dependent component of this drug contributes to the high barrier to MZA resistance by *M. tuberculosis*.

After confirming resistance phenotypes, six independently selected isolates of H37Rv Δ*pncA* and four of BCG were analyzed using whole genome sequencing. All ten isolates had single nucleotide mutations within the promoter region of *mscR* (Figure 3D) encoding a mycothiol oxidoreductase which is critical for formaldehyde tolerance of mycobacteria (20). To confirm the protective nature of mutations in the *mscR* promoter region against aldehyde stress, we exposed wild-type and MZA resistant mutants to methenamine, a bactericidal antibiotic that acts through the spontaneous release of formaldehyde and is commonly used to treat urinary tract infections (21). While the MIC of methenamine for the wild-type BCG strain was 900 µM, the MZA resistant strains showed resistance to 7.1 mM methenamine (Figure 3E,F).

To confirm whether the identified promoter mutations conferred MZA resistance through increased MscR expression, we constructed GFP reporter fusions to the *mscR* promoters from wild-type and C-15T resistant variant in the integrative reporter plasmid pYUB1227. Reporter constructs were integrated at the L5 attachment site in BCG and confirmed strains were grown to exponential phase. After fluorescence was normalized to the OD_600_ a 1.7 fold increase in fluorescence was observed for the *mscR* promoter variant relative to the wild-type *mscR* promoter (Figure 3G) confirming that resistance was due to increased expression of mycothiol oxidoreductase. To validate the role of *mscR* in intrinsic resistance of *M. tuberculosis* to MZA we constructed a H37Rv Δ*mscR* strain and assessed MZA susceptibility. This strain showed an eight-fold enhanced level of susceptibility relative to the parental control (Figure 3H).

Given the role of MscR in MZA susceptibility and resistance, we reasoned that cysteine supplementation may antagonize MZA action through interaction with thiol metabolism. To test this hypothesis, BCG wild-type and *mscR* promoter variants were exposed to inhibitory concentrations of 900 µM methenamine or MZA in the presence of varying concentrations of cystine as a source of cysteine (Figure 3I,J). For wild-type BCG, 4.1 mM cysteine supplementation was highly antagonistic for both MZA and methenamine. For the *mscR* promoter mutant, cysteine supplementation was highly antagonistic for MZA at an eight-fold lower concentration than was required for the wild-type strain. Together, these data demonstrate a central role for thiol metabolism in MZA action and establish increased MscR expression as a mechanism for MZA and methenamine resistance.

### Genome-wide fitness profiling and chemical biology reveal the unique self-potentiating mechanism of MZA

To better understand pathways that modulate resistance and susceptibility of *M. tuberculosis* to PZA, POA and MZA, we conducted fitness profiling by using Tn-seq. A saturated library of ∼70,000 transposon mutagenized H37Ra was prepared using the *himar1* mariner transposon (22). Colonies were harvested and homogenized to form a pooled library and aliquots were precultured in the absence of drug. Exponentially growing libraries were then subcultured and grown for 5 generations in the absence of drug or in the presence of a concentration of PZA (233 µM), POA (105 µM) or MZA (112 µM) that resulted in a two-fold extension of the generation time. These concentrations and number of generations were chosen to allow for the highest resolution of gain- and loss-of-fitness conferred by transposon insertion. Following growth, cells were harvested, genomic DNA was extracted and deep-sequencing of transposon junctions was performed to determine relative abundance of each insertion (8). Intra-population fitness was determined by comparing abundance of individual insertion sites before and after enrichment using a previously established methodology as described in Opijnen et al. 2009 (23). To determine the change in fitness of specific Tn-insertion mutants, the relative fitness of each insertion site was evaluated by calculating the change in fitness with drug compared to the absence of drug. Insertions with relative fitness scores > 1 were considered gain-of-fitness mutants, while relative fitness scores < 1 were considered loss-of-fitness mutants. Consistent with our previous findings on PZA and POA susceptibility (8), insertions in genes involved in phosphate utilization and transport (*pstA1, pstC2, pstS3, phoT*), cell wall stress response regulator (*sigE*), and acyl-CoA synthetase (*fadD31*) demonstrated substantially greater fitness in the presence of all 3 drugs (Figure 4A-C). We also observed a consistent gain-of-fitness for mutants with insertions in the PDIM locus which correlates with our RNA-seq data and previous indications that PDIM mutants are moderately resistant to PZA (19). In addition, *pncA* insertions were found to have a gain-of-fitness in PZA and MZA treated samples, but not POA treated samples, supporting our observation that PncA plays a supporting role in MZA activity (Figure 4A-C). We previously described loss-of-function mutations in *fadD2* that enhanced susceptibility to PZA (24), yet, such mutants could not be analyzed in this screen due to lack of abundance of respective insertions in our library. Importantly, insertions in the *mscR* locus harbored substantial fitness defects in MZA treated samples relative to PZA and POA treated samples (Figure 4C). This finding is consistent with our observation that MZA susceptibility is modulated by MscR activity in mycobacteria. Interestingly, strains with insertions in *sseA*, recently shown to be critical for maintaining thiol homeostasis in *M. tuberculosis* (25), showed a substantial loss-of-fitness in the presence of PZA, POA and MZA, consistent with a direct connection between thiol metabolism and activity of these drugs. Collectively these data indicate MZA, PZA and POA show conserved interactions within *M. tuberculosis* metabolism, yet, that MZA action is distinct due to its unique ability to interfere with thiol homeostasis.

**Figure 4.**
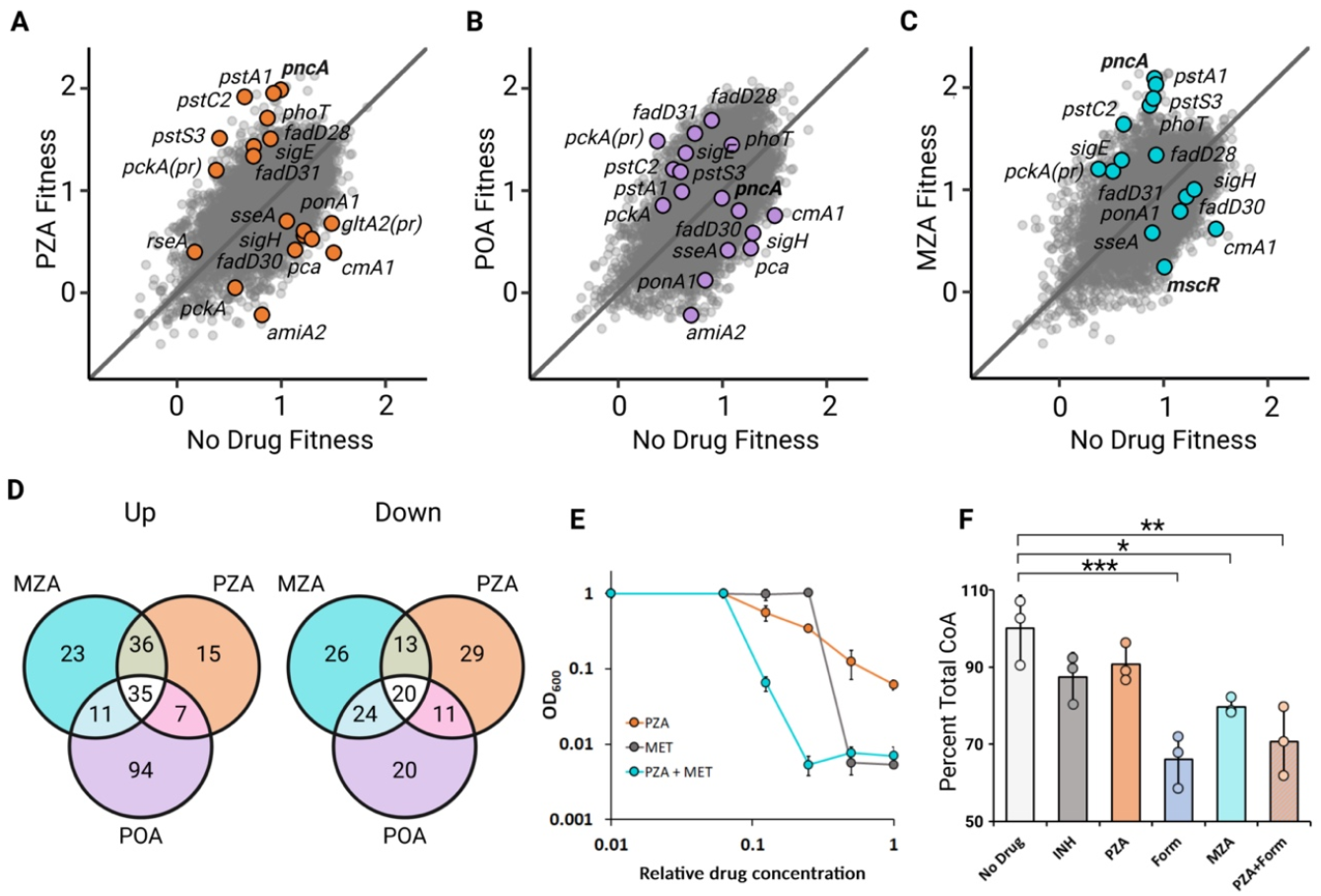
Genome-wide fitness profiling of *M. tuberculosis* transposon insertion mutants and chemical biology reveal the unique self-potentiating mechanism of MZA. Saturated library of Tn-insertion mutants treated with **A**, 233 µM PZA **B**, 105 µM POA and **C**, MZA 112 µM MZA at pH 6 to achieve a 50% reduction in growth rate. Each point corresponds to a specific TA site harboring a transposon insertion within the genome. The plot shows every transposon insertion identified. Highly depleted or enriched TA sites are highlighted with genes of interest noted. **D**, Venn diagram showing genes enriched for (left) and depleted (right) shared by all three treatments. **E**, DiaMOND assay showing exposure of *M. tuberculosis* H37Ra to a geometric series of PZA and methenamine (MET) alone and in combination. **F**, Abundance of CoA from *M. tuberculosis* H37Ra at pH 6.0 treated with DMSO (No Drug), INH, PZA, formaldehyde (Form), MZA and a combination of PZA and formaldehyde (PZA+Form). Values were normalized to total protein and presented as percentages relative to the no drug control. Significance was determined using a one-tailed Dunnett’s test with Bonferroni correction. All assays were performed in biological triplicate with mean displayed and error-bars indicative of standard deviation.

Since MZA action shares many features with PZA but does not require exposure to low pH for this aspect of its activity, we next evaluated whether aldehyde can potentiate PZA action *in lieu* of exposure to low pH. To do so, we conducted a DiaMOND assay (26) testing for synergy between methenamine and PZA at circumneutral pH (Figure 4E), as well as methenamine and RIF, INH, EMB and POA (Figure S2). While the individual MIC of PZA at neutral pH surpassed 8 mM and MIC of methenamine was 440 µM, the combination of the two produced a Fractional Inhibitory Concentration Index (based on the IC_90_ of individual drugs) (FICI) of 0.375 indicative of synergy (26), indicating that aldehyde stress sufficiently potentiates PZA activity (Figure 4E). Yet, in the case of other drugs, the interaction was found to be additive (Figure S2).

To further characterize the synergistic dual action of MZA, we assessed whether MZA could participate in depletion of CoA abundance (27). To test this hypothesis, we measured intracellular CoA abundance using H37Ra treated for 8 hours with formaldehyde, PZA, MZA, or a combination of PZA and formaldehyde. INH was included as a control. Previous reports demonstrate a 50% reduction in CoA abundance after 24 hours of PZA exposure (19, 24). With the shorter exposure time use here, reduction of CoA in PZA treated cells was mild and did not meet a significance threshold (Figure 4F). Unexpectedly, INH was also found to produce a similar mild decrease in CoA abundance that also did not meet a significance threshold. In contrast, CoA reduction in cells treated with MZA or formaldehyde with or without PZA resulted in a 20-30% reduction in CoA abundance (Figure 4F), indicating that aldehyde drives a more rapid depletion of CoA than does PZA.

### MZA shows superior potency relative to PZA against intracellular *M. tuberculosis*

Given the critical role of cell-mediated responses in PZA efficacy and the growing problem of drug-resistant *M. tuberculosis*, we assessed the ability of MZA to kill intracellular *M. tuberculosis* in resting or IFN-γ activated macrophages. RAW 264.7 macrophages were exposed to *M. tuberculosis* H37Rv or H37Rv Δ*pncA* at a multiplicity of infection of 1:1 and extracellular bacteria were removed by washing adhered macrophages. For assessing the impact of macrophage activation on MZA efficacy, 5 ng/ml IFN-γ was added one day prior to *M. tuberculosis* infection and replenished every other day. MZA and PZA were used at a concentration of 900 µM and compared to treatment with no drug (DMSO). At the concentration tested, PZA showed limited activity against H37Rv and no activity against H37Rv Δ*pncA* in resting and activated macrophages (Figure 5A-D). In contrast, MZA caused a greater than 3 log_10_ decrease in H37Rv viability by day four of treatment in both resting and activated macrophages, and after a week, bacterial loads fell below the detection threshold (Figure 5A,B). Comparable findings were also observed when macrophages were infected with H37Rv Δ*pncA* (Figure 5C,D), indicating that the primary bactericidal action of MZA against intrabacterial bacilli is mediated through its aldehyde release mechanism. Taken together these data demonstrate that the robust bactericidal activity of MZA is both independent of macrophage potentiation and highly effective against intracellular PZA resistant isolates.

**Figure 5.**
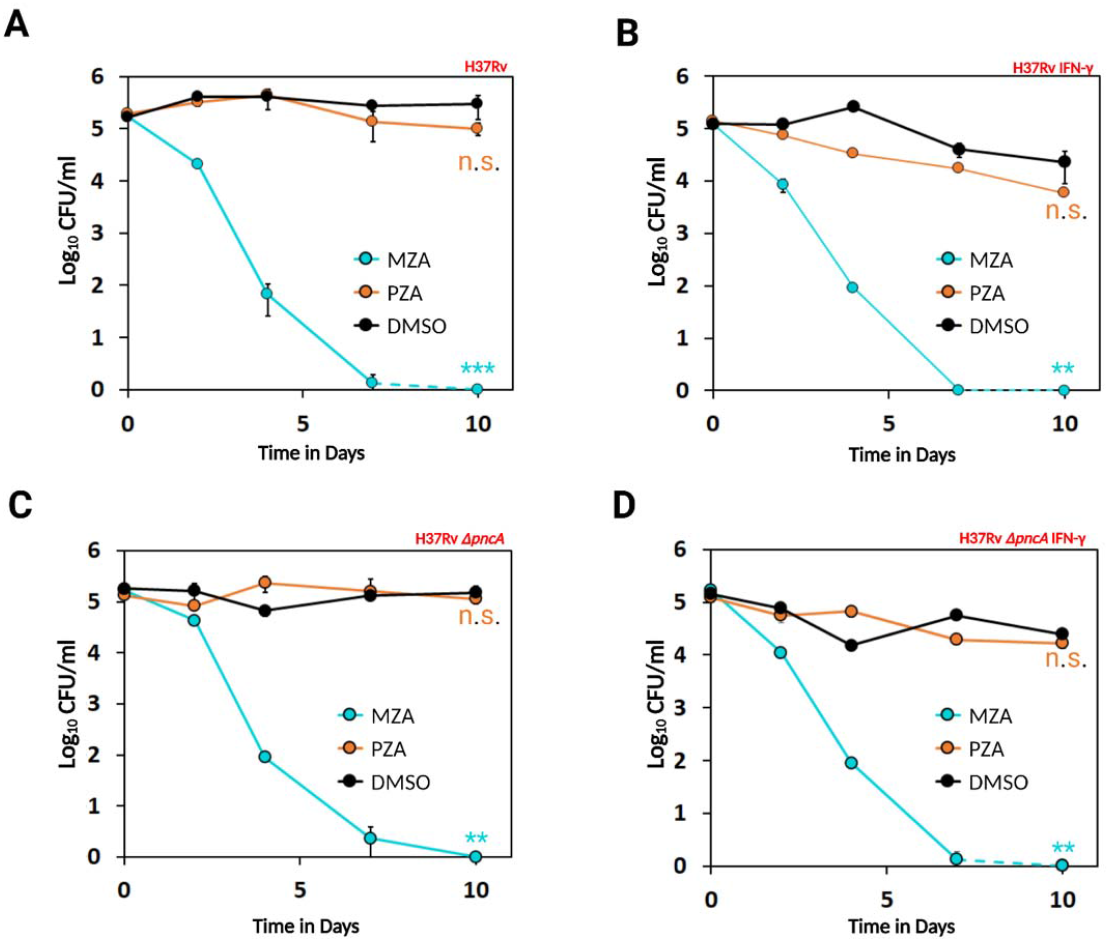
MZA efficacy against PZA susceptible and PZA resistant *M. tuberculosis* strains in resting and activated macrophages. **A**, CFU comparison between RAW 264.7 macrophages infected with H37Rv and treated with either MZA, PZA or DMSO. **B**, H37Rv infected macrophages were activated using IFN-γ. **C**, Same experimental design as in A but using the PZA resistant H37Rv Δ*pncA* strain to test efficacy of MZA against PZA resistant strains. **D**, RAW 264.7 macrophages activated with IFN-γ and infected with PZA resistant H37Rv Δ*pncA*. Cells were treated with 0.9 mM MZA or PZA with daily media exchange. All assays were performed in biological triplicate with mean displayed and error-bars indicative of standard deviation. Significance was determined via Kruskal-Wallis test combined with post hoc Dunn test with *p*<0.01 (_**_), *p*<0.001 (_***_).

## Discussion

In this study we employed a combination of approaches in functional genomics, bacterial physiology and chemical biology to characterize the synergistic dual-action *M. tuberculosis*-specific mechanism of the PZA analog MZA. Our analyses revealed an overlap in a central component of the mechanisms of action of PZA and MZA while also resolving the underlying basis for differences in their antitubercular activity. These findings led us to identify a self-potentiating mode of action specific to MZA that is both unique from and more potently bactericidal than its PZA counterpart. The unique features of MZA, which include rapid killing, high target specificity, low rates of resistance and independence from environmentally-driven potentiation, can be attributed to the drug-specific release of aldehyde resulting in disruption of thiol homeostasis. The rate of aldehyde release, as determined by mass spectrometry, takes place over the course of hours, which allows for elimination of bacilli from infected macrophages within a matter of days. The superior feature of MZA is orchestrated by the synergistic actions of its two components, aldehyde release followed by PZA collateral susceptibility (Figure 6). The rapid engagement of aldehyde elicits bactericidal activity against both actively growing and dormant cells alike while simultaneously sensitizing any survivors to the activity of PZA. This potentiation event likely ensues via depletion of biologically active thiols, resulting in bacterial growth arrest and increased demand for CoA with consequent disruption of fatty acid biosynthesis, a pathway known to be perturbed by PZA (15, 28). The mechanism proposed here is further confirmed by synergistic activity of methenamine with PZA and the antagonistic activity of cysteine against MZA.

**Figure 6.**
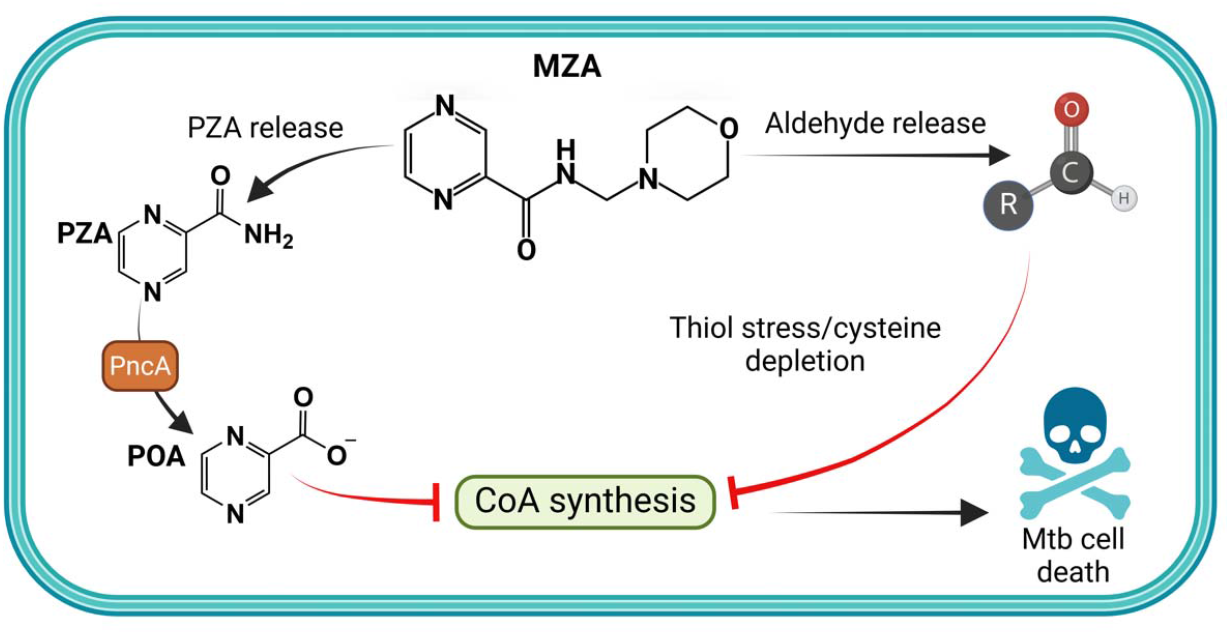
Dual Action Mechanism of Morphazinamide. MZA releases aldehyde and PZA in *M. tuberculosis*. PZA is converted to POA which interferes with CoA synthesis. Aldehyde release drives thiol stress resulting in synergistic disruption of CoA metabolism with PZA and rapid bacterial cell death.

The robust activity of MZA against non-growing *M. tuberculosis* suggests that entry into the cytoplasm is likely a passive event. Hence, MZA is effective at killing *M. tuberculosis* independent of metabolic activity or growth state of the cells. Passive entry of MZA into the cytoplasm also supports a low likelihood for emergence of drug-resistant strains via mutations that alter drug transport. While low-level resistance to aldehyde release can be achieved via activation mutations in the *mscR* promoter, achieving full resistance to MZA will likely be a challenge due to the promiscuous nature of aldehyde-mediated damage. Moreover, we find that PZA resistant strains of *M. tuberculosis* and *M. bovis* BCG with *mscR* promoter-up mutations still show measurable susceptibility to MZA, suggesting either that the rate of internal aldehyde release is greater than the MscR turnover rate or that other aldehydes that cannot be effectively neutralized by MscR are released by MZA. In either case, the bacilli will be faced with a greater adaptive burden to develop full resistance to MZA (Figure 6).

Given the pivotal role of PZA in TB treatment regimens and the limited activity of this drug in models of compromised immunity (11), it will be critical to identify PZA analogs whose activity is not dependent upon cell-mediated responses. In the present study, aldehyde release from MZA was able to reduce bacterial loads in resting macrophages to levels below the limit of detection in under a week. Based on loss of PZA efficacy in the athymic mouse model (11), it has been speculated that antitubercular activity of PZA may be diminished in the context of T cell deficiency or dysfunction due to insufficient macrophage activation, resulting in reduced treatment efficacy and higher rates of drug-resistance (9). Circumnavigating the need for host-driven potentiation of PZA may prove to be a valuable approach in the treatment of *M. tuberculosis* in the context of compromised immunity.

Interestingly, it has recently been recognized that production of host-derived aldehydes are modulated via IFN-γ activation and play a role as antimicrobial effectors against intracellular pathogens, such as *M. tuberculosis* and *Francisella tularensis* (29, 30). Whether host-derived aldehydes participate in the bactericidal activity of PZA *in vivo* has yet to be determined, but, may represent an opportunity for host-directed therapy for bolstering the action of this critical antitubercular agent. Further, microbe-derived aldehyde production may also represent an opportunity that can be explored for novel therapeutic discovery. For example, selective disruption of cytokinin metabolism has been shown to result in accumulation of toxic aldehyde species in *M. tuberculosis* and further sensitize the bacilli to other host stressors (31). Further, as noted by Darwin and Stanley (29), mycobacterial glycolysis may represent an exploitable target for driving toxic aldehyde accumulation in *M. tuberculosis* (32). Along these lines, it is curious to note that strains bearing loss-of-function mutations in *glpK* (encoding glycerol kinase) show a selective advantage both in humans and in mice treated with antitubercular agents (33, 34). Since disruption of glycerol phosphate synthesis would reduce the level of endogenous metabolic aldehydes through limitation of glyceraldehyde phosphate and dihydroxyacetone phosphate synthesis, glycerol kinase likely represents a key mediator in endogenous mycobacterial aldehyde production (29). In support of this concept, Dick and colleagues previously determined that mutations in *glpK* are associated with PZA resistance *in vitro* (19). Our findings, coupled with the emerging role of host- and microbe-derived aldehydes in compromising the fitness of *M. tuberculosis*, advocate for further evaluation of means for targeted delivery of aldehyde-releasing antimicrobial agents, such as MZA, for development of improved therapeutic approaches.

## Materials and Methods

### Standard growth conditions

Middlebrook 7H9 broth (Difco) supplemented with 10% Middlebrook OADC (Difco), 0.02% (V/V) glycerol and 0.05% (V/V) tyloxapol were used for all experiments involving liquid *M. tuberculosis* and *M. bovis* cultures. Depending on experimental requirements, the media were adjusted to pH 7.2 or pH 5.8. Cultures were incubated shaking at 100 RPM 37ºC. All solid media used in experiments involving *M. tuberculosis* and *M. bovis* were based on Middlebrook 7H10 (Difco) supplemented with 10% Middlebrook OADC, 0.02% (V/V) glycerol. For *M. smegmatis* the same media were used with the exclusion of OADC and replacing glycerol with 0.2% (W/V) dextrose. ESKAPE pathogens and *E. coli* (for plasmid amplification) were propagated in Luria-Bertani broth (LB, Difco) under standard growth conditions (37°C shaking at 250 RPM).

### Strain construction

The knockout strains (H37Rv Δ*pncA* and H37Rv Δ*mscR*) and were constructed using the ORBIT system (35). H37Rv was electroporated with pKM461. After selection on 50 µg/mL kanamycin, H37Rv pKM461 was induced with anhydrotetracycline and electroporated with pKM464 and the position specific *pncA* oligonucleotide 5’-TACCTCGGCGCCACGGCGGCGGACCCGGCCCGCGCCCGGTGGCTCCT GCACTTCGGCATGGTGGGCCGCAGGTTTGTACCGTACACCACTGAGACCGCGGTGGTTGA CCAGACAAACCCCTCGACTCGCTTCCGACAGCACCTCGAAGACCGCTTCGGGTGCGTGAG CACGCTGGGCGGTTCGCAGTG-3’ or *mscR* 5’-GCGCGCATGGTCAGCGACGCTACACCGTAGGTTGGACACCATGAGTCAGACGGTGCGCGGTGTGATCGCAGGTTTGTCTGGTCAACCACCGCGGTCTCAGTGGTGTACGGTACAAACCAAGGTATTGCGTTCGGTGGTGATGTTGTGATGGCCGCCATCGAGCGCGTCATCACCCACGGCACCTTCGA-3’. Transformed bacteria were then plated on supplemented 7H10 medium containing 50 µg/mL hygromycin B and 10% sucrose to select for recombinants and counter select against pKM461. Individual colonies were then re-streaked, confirmed via PCR, and full genome sequenced to validate desired modification and no additional mutations.

Fluorescent reporter *mscR* strains were constructed using a modified integrative reporter plasmid pYUB1227. Restriction enzymes *Pvu*I and *Mfe*I (NEB) were used to prepare the backbone and the prompter inserts, followed by a T4 ligation. Primers used to validate promoter changes were mscRF 5’-TTGCAACGCATCCCTGATCT-3’ and mscRR 5’-AGGGCAGATTGTGTGGACAG-3’. Additional validation was performed via Nanopore sequencing. Plasmid amplification was performed in *E. coli* DH5α. Plasmids were purified using a QIAprep Spin Miniprep Kit (cat.# 27104). BCG Pasteur 1173P2 was then transformed using previously prepared plasmids via standard electroporation protocol. Successful transformants were selected for on standard media amended with 50 µg/mL hygromycin B and validated via genomic DNA extraction and PCR using validation primers described above.

### Determination of minimum inhibitory concentrations

Bacteria were grown in 30 mL PETG square media bottles (Nalgene) containing 5 mL of 7H9 media under standard growth conditions (37°C shaking at 100 RPM) to OD_600_ of 0.15-0.25. Bacteria were then diluted to OD_600_ of 0.01 and allowed to incubate for 2 weeks in presence of the compound(s) being tested. Concentrations of compounds were determined as a function of a geometric series (0.05, 0.1, 0.2,0.4, 0.81, 1.62, 3.25 and 6.5 mM) for PZA, (0.06, 0.11, 0.22, 0.45 and 0.9 mM) for MZA and (0.06, 0.11, 0.22, 0.44, 0.88, 1.75 and 3.5 mM) for methenamine. MIC values were then calculated by plotting inhibition curve and calculating based on the appropriate slope intercept.

### Antagonism assays

Bacteria were prepared in the same way as for MIC assays and were then inoculated into 7H9 media containing either 0.9 mM methenamine, 0.9 mM MZA or DMSO (vehicle control) and antagonist (pantothenate or cystine) at indicated concentrations. Pantothenate was used at a single concentration of 230 µM, cystine concentrations ranged from 15 µg/mL to 500 µg/mL in 2-fold increments. The media pH was adjusted to 7.2 and cells were incubated in 96-well plated for 8 days at 37°C with no shaking. Measurements were performed using a BioTek Synergy H1 plate reader (Agilent). All experiments were performed in triplicate.

### Bacterial kill curves

To assess the bactericidal activity of MZA and methenamine, against exponentially growing cells, bacteria were grown in 30 mL bottles containing 5 mL of 7H9 media under standard growth conditions (37ºC shaking at 100 RPM) to OD_600_ of 0.15-0.25. Bacteria were then diluted to OD_600_ of 0.03 (approximately 5X10^6^ bacilli) and plated on 7H10 media to determine the starting population of bacteria. After addition of (0, 0.28, 0.41, 0.61 or 0.9 mM of MZA) and (0.06, 0.11, 0.22, 0.44, 0.88, 1.75 and 3.5 mM) µg/mL Methenamine, CFU counts were determined at 0, 2-, 4-, 7- and 14-day time intervals.

To assess bactericidal activity of PZA and MZA against nutrient limited *M. tuberculosis*, bacteria were grown in 30 mL bottles containing 5 mL of 7H9 media under standard growth conditions (37° C shaking at 100 RPM) to OD_600_ of 0.15-0.25. Bacteria were then washed three times with 1xPBS saline to remove any residual media and resuspended in 1xPBS saline amended with 0.05% tyloxapol. Prior to addition of the drugs CFU/mL were determined by serial dilution plating on 7H10 media, 900 µM PZA, MZA or DMSO were then added to the buffer and CFU measurements were taken at predetermined intervals 0, 2-, 3-, 4-, 7- and every 7 days afterwards until 62-day timepoint was reached. For MBC measurements under starvation conditions with multiple treatments, bacteria were prepared in the same way as for a single treatment. However, at every timepoint culture was washed three times to remove residual antibiotics and split into two cultures. One of the two cultures were treated with DMSO while the other was amended with fresh MZA for a final concentration of 900 µM.

### RNA-seq

For bulk RNA-seq library preparation and sequencing, exponentially growing H37Rv and H37Rv Δ*pncA* were grown to the OD_600_ of 0.15 and inoculated with 200 µM MZA, PZA or equivalent volume of DMSO (as a vehicle control). Cells were then incubated under standard condition for 24 hours prior to harvesting. Following incubation, TRIzol RNA purification was performed. Cells were chilled, pelleted and resuspended in a 500 µL of TriReagent (Invitrogen) with 1% polyacryl carrier (Molecular Research Center) and lysed using 250 µL of 0.1 mm zirconia beads (BioSpec). Samples were then centrifuged and supernatant was transferred to a clean 2 mL microcentrifuge tube and combined with 50 µL of 5-bromo-3-chloro-propane. Following a 10-minute incubation samples were centrifuged aqueous phase was removed and transferred to a clean 1.5 mL Eppendorf tube where samples were treated with 250 µL of isopropanol. Following a 10-minute incubation samples were centrifuged again, isopropanol was removed and RNA coating the tube was washed with 300 uL of 75% EtOH. After the final wash, RNA was dried and eventually resuspended in 50 uL of DEPC-treated water (Invitrogen). A DNA-free ^(™)^ DNAse treatment (Invitrogen) was used to remove residual DNA from the samples prior to sequencing. Samples were treated with Ribo-Zero Plus kit and Libraries were prepared using an Illumina Stranded Total RNA Prep Ligation. Sequencing was performed at the University of Minnesota Genomic Center using a 50 paired-end NovaSeq S-prime platform.

For bulk RNA-seq analysis, the pipeline used for preprocessing raw fastq files can be found at https://github.com/MDHowe4/RNAseq-Pipeline. Quality control of RNA sequencing read quality was assessed with FastQC. Read length thresholding and t-overhang trimming of forward and reverse reads was completed with Cutadapt with minimum read length cutoff of 30 bp. Reads were mapped to the M. tuberculosis H37Rv reference genome (NC_000962.3) using the STAR aligner without spliced alignment detection (--alignIntronMax 1) (36). Total reads per gene were counted using featureCounts (37). Genes with <10 reads mapped occurrences across wild-type and mutant experiments were not included in further analysis. Differentially expressed genes (DEGS) were found using the negative binomial generalized linear model of DESeq2 (38). Genes were considered differentially expressed if they displayed a log_2_ fold-change ≥ 1.5 or ≤ -1.5 and adjusted *P*-value ≤ 1x10^-6^. Volcano plots were generated using ggplot2 (39).

### Mass spectrometry

For mass spectrometry analyte preparation, for each sample, a total of 40 mL of bacteria were grown under standard growth conditions at pH 5.8. Upon reaching OD_600_ of 0.4, bacteria were inoculated with either 250 µM MZA or PZA. After three days of treatment the samples were pelleted. Cell extract samples were prepared for mass spectrometry analysis by resuspending pellets in 1 mL of extraction buffer (40% methanol, 40% acetonitrile and 20% water) and bead beaten using 250 µL of 0.1 mm zirconia beads (BioSpec). Samples were then centrifuged to remove the solids. Liquid fractions were then filtered using 22 µm filter. A portion of the filtrate was then used to determine protein content using a Pierce^TM^ BCA Protein Assay Kit (Thermo Scientific). The remaining samples were further purified using a 3 kDa column (PALL Life Sciences). Supernatant sample preparation was performed in the identical way as cell extract, except for the omission of the first bead beating step. Bacterial cell extracts were diluted 10-fold (10 µL in 100 µL) in DI water. 100 µL of diluted solution was treated with 100 µL methanol: water solution containing *para*-aminosalicylic acid (PAS, internal standard), mixed well by vortexing, centrifuged (2 min, 15000 × g), and supernatant (150 µL) was analyzed by LC-MS/MS.

For LC-MS/MS analysis, reverse-phase LC was performed on a Zorbax Eclipse XDB-C8 column (150 mm, 4.6 mm, 5 µm; Agilent) using Shimadzu UFLC XR instrument. The elution gradient was carried out with binary solvent system consisting of 0.1% formic acid in H_2_O (solvent A) and 0.1% formic acid in MeCN (solvent B). A linear gradient profile with the following proportions (v/v) of solvent B was applied (t (min), %B): (0, 5), (1, 5), (4, 95), (6, 95), (7, 5), (8, 5) with 2 min for re-equilibration to provide a total run time of 10 min. The flow rate was 0.5 mL/min and the column oven was maintained at 40°C. The injection volume was 10 µL. The retention times for MZA, PZA, POA and PAS (internal standard) were 3.5, 2.6, 3.8 and 6.8 min, respectively. MS/MS analysis were carried out using triple quadrupole/linear ion trap instrument (AB SCIEX QTRAP 5500). MZA, PZA and POA peak areas were calculated using MultiQuant, version 2.0.2. MZA, PZA and POA areas were normalized to p-amino salicylate areas and the MZA, PZA and POA concentration in samples were determined using the standard curves of MZA, PZA and POA. For the preparation of MZA, PZA and POA standard curve, average area of standard 0 was subtracted from MZA, PZA and POA of all higher standards prior to area normalization. Sample concentrations were back calculated to account for dilutions. Each sample was performed in triplicate.

### Transposon mutagenesis and transposon sequencing

*M. tuberculosis* H37Ra was mutagenized with the mariner *himar1* transposon using the phAE180 temperature-sensitive mycobacteriophage (40). Approximately 70,000 independent mutants were selected on 7H10 media amended with 50 µg/mL kanamycin. Transposon mutants were pooled, homogenized, then aliquoted into 25% (V/V) glycerol stocks and stored at -80°C. Growth curves to establish 50% inhibition of growth rate using a geometric series of drug concentrations were carried out using 7H9 medium at a pH of 6.0. Cultures were seeded at OD_600_ of 0.01 in 80 mL of medium using transposon mutants pre-cultured in a shaking incubator for 48 hours at 37°C. Enrichment was carried out in the presence or absence of drugs for 5 generations as determined by OD_600_. After 5 generations, cultures were flash frozen in liquid nitrogen for genomic DNA (gDNA) extraction. gDNA extraction was carried out using a previously established protocol (41). Library prep and sequencing was performed at the University of Minnesota Genomics Center with read mapping performed using a previously established protocol (8). DNA fragmentation and Illumina P7 (CAAGCAGAAGACGGCATACGAGAT) ligation was performed. Amplification of transposon site junction was carried out using a P7 (CAAGCAGAAGACGGCATACGAGAT) and mariner-specific Mariner_1R_TnSeq_noMm primer (TCGTCGGCAGCGTCAGATGTGTATAAGAGACAGCCGGGGACTTATCAGCCAACC).

Amplification of himar1-enriched samples was performed using a P5 indexing primer (AATGATACGGCGACCACCGAGATCTACAC[i5]TCGTCGGCAGCGTC; [i5] barcode sequence) and a P7 primer HotStarTaq master mix kit (Qiagen) after a 1:50 dilution. Sequencing was performed on an Illumina NovaSeq. The 5’ ends of reads were trimmed to remove adapter and transposon sequence using Cutadapt to leave the TA-gDNA junction intact (42). Trimmed reads with less than 18 bp were discarded and reads were mapped to the M. tuberculosis H37Ra genome (GenBank Accession: CP000611) with 1 bp permitted mismatch using Bowtie 2 (43). Mapped reads were printed in SAM format and counted per TA dinucleotide sight using SAMreader_TA script (44). Scripts used for read processing may be found on GitHub (https://github.com/MDHowe4/Himar1-TnSeq-Pipeline).

After using the TRANSIT hidden Markov model to obtain number of insertions at genome positions (45), the relative abundance of insertions at TA sites was calculated for each sample. TA sites lacking mapped insertions at the start of enrichment were omitted. TA sites without mapped insertions at the end of enrichment that were present at the start of the enrichment were set to the limit of detection without contributing to the total number of insertions. Relative fitness of site-specific mutants were calculated with a previously characterized function using the population expansion factor, as well as the relative number of insertions at the beginning and end of enrichment (23).

### Isolation of spontaneous mutants

To isolate spontaneously MZA resistant mutants, H37Rv Δ*pncA* and BCG cells were grown up to mid-log phase for selection. 1 mL of culture diluted to OD_600_ of 0.3 was plated onto standard 7H10 agar supplemented with 409 µM MZA for H37Rv Δ*pncA*, and 607 µM for BCG. Plates were then incubated at 37°C in stationary incubator for 14-21 days. Resistant mutants were then picked and streaked again for isogenic backgrounds, then grown again for 14-21 days at 37°C in a stationary incubator. Isogenic colonies were picked, then grown in standard 7H9 broth for 14 days followed by genomic DNA extraction using a the previously described DNA extraction methodology. Illumina sequencing was performed at SeqCenter. Read alignment and variant calling to the H37Rv reference genome (NC_000962.3) and BCG reference genome (BCG Pasteur 1173P2) was performed using Breseq version 0.36.0.

### Coenzyme A detection assays

Bacteria were prepared for extraction in the same way as described in mass-spectrometry methods. 80 mL per treatment sample of H37Ra was grown to an OD_600_ of 0.13 in 7H9 media (pH 6.0) in the shaking incubator at 100 RPM and 37°C. All of the samples were then treated with 200 µM of PZA, MZA, formaldehyde, formaldehyde and PZA, or 0.292 µM isoniazid. After an eight-hour exposure cells were harvested and resuspended in 400 µL of ice cold 1xPBS and lysed using zirconia beads. Samples were then centrifuged and filtered using 0.22-µm filter. 50 µl aliquots were then taken to perform a BCA assay (Pierce™ BCA Protein Assay Kit, ThermoScientific). The remaining extracts were then filtered one more time using a using a 3 kDa column (PALL Life Sciences). Twice filtered lysate was then used to perform fluorometric assay using the Free Coenzyme A Assay kit (Sigma-Aldrich, cat# MAK034 or MAK504). The readout was then normalized to the protein abundance, then coenzyme A levels of each sample were compared to the mean DMSO control for interexperimental comparison. All assays were performed in triplicate.

### Macrophage Infections

RAW 264.7 macrophages were prepared by seeding density of 1.0-2.0 x 10^5^ cells per well in DMEM/F-12 medium containing 10% FBS without antibiotics (DMEM/F-12 complete) in 12 well (1 mL/well) plates. In order to allow for cells to adhere, macrophages were incubated overnight in a humidified 5% CO_2_ chamber. The following day cells were rinsed with Hank’s buffer three times and replenished with fresh DMEM/F-12 media. A 5 ng/mL of IFN-γ was added to the well containing macrophages meant to be activated. Following a 14-16h incubation, macrophages were washed again in Hank’s buffer and replenished with fresh, drug and interferon free DMEM media. Upon transfer of macrophages into BSL-3 space, media was removed, cells were washed with Hank’s buffer three times and fresh DMEM media containing *M. tuberculosis* H37Rv. An MOI of 1:1 was used for the initial infection. To prepare bacteria, H37Rv was grown under standard growth conditions. Upon reaching OD_600_ of 0.25, culture was washed three times, resuspended, and then diluted in DMEM media to the 1.0-2.0 x 10^5^ CFU/mL. Following a 2h infection, infected macrophages were washed three times with Hank’s buffer and resuspended in complete DMEM media amended with 900 µM MZA, PZA or DMSO (as a vehicle control) as well as 5ng/ml of IFN-γ for activated macrophages. Media was changed daily for the duration of the experiment with re-induction via IFN-γ taking place on every other day. During every media change, cells were washed three times with Hank’s buffer and adherence and appearance of macrophages was monitored via a stereoscope. On plating days, cells were washed as described above, and macrophages were lysed via TritonX-100. The lysate was then serially diluted in 0.05% Tyloxapol 1xPBS pH 7.2 buffer using 10-fold dilution in a 96well plate format and plated on 7H10 media. Plates were incubated for two weeks and colonies counted for CFU determination. All experiments were performed in triplicate.

### MZA synthesis and NMR analysis

Formaldehyde 37% in water (89 µL, 1.9 mmol) was prepared with PZA (200 mg, 1.6 mmol) and morpholine (1.1 mL, 12.9 mmol) in 100% ethanol solvent with one drop of concentrated HCl. The reaction was refluxed for 6 hours and neutralized with one drop of 1 N NaOH. The crude mixture was evaporated and separated via silica gel flash chromatography with methanol/dichloromethane solvent system. The pure compound (>95%) was a white solid. The calculated yield was 63.7%. Observed (M+H) ^+^ of 223.0 m/z with mass spectrometry analysis.

^1^H NMR (400 MHz, CDCl_3_-d, ppm): δ = 9.35 (s, 1), 8.71 (d, 1), 8.48 (d,1), 8.12 (s, 1), 4.29 (d, 2), 3.65 (t, 4), and 2.58 (t, 4). ^13^C NMR (100 MHz, DMSO-d_6_ ppm): δ = 164.2, 148.0, 145.0, 144.1, 143.7, 66.4, 60.8, and 50.5. Purity: >99%.

### Quantification and statistical analyses

Where applicable, 95% confidence intervals were used to determine statistical significance. Numbers of biological replicates and statistical tests that were employed are described in respective figure legends.

## Materials availability

Materials will be made available upon reasonable request and may require completion of a Material Transfer Agreement and payment for related expenses.

## Data and code availability

All data are available in the main manuscript or supplemental information. Primary sequencing data from RNA-seq, Tn-seq, and whole-genome sequencing studies are publicly available through the National Center for Biotechnology Information via SRA link https://www.ncbi.nlm.nih.gov/sra/PRJNA1104292.

Code for bulk RNA-seq analysis pipeline used for preprocessing raw fastq files can be found on GitHub at https://github.com/MDHowe4/RNAseq-Pipeline. Code used for read processing can be found on GitHub https://github.com/MDHowe4/Himar1-TnSeq-Pipeline.

## Acknowledgments

We would like to acknowledge the following individuals and institutions for their invaluable contributions to this work. We express our gratitude to the NIH for their generous grant support Grant Numbers: R01 AI123146 (A.D.B), T32 GM008244 (S.V.) and T32 HL007741 (L.O. and T.A.C.). We also thank the University of Minnesota Dinnaken Fellowship for support of T.A.C. We are also grateful to the University of Minnesota Genomics Center and University of Minnesota Biosafety Level 3 Program Core Facilities, and SeqCenter for their technical support and expertise. We also would like to acknowledge Biorender which was used for the preparation of the figures. Lastly, we would like to thank Dr. Elise Lamont for her input and support for macrophage work and Dr. Subhankar Panda for determination of MZA purity.

## Notes

https://www.ncbi.nlm.nih.gov/sra/PRJNA1104292

https://github.com/MDHowe4/RNAseq-Pipeline

https://github.com/MDHowe4/Himar1-TnSeq-Pipeline

### Competing Interest Statement

The authors have declared no competing interest.

### Summary of Updates

This manuscript has been updated based on initial reviews by other experts in the field.

