## Supplemental Materials for "Mechanism of the Dual Action Self-Potentiating Antitubercular Drug Morphazinamide"

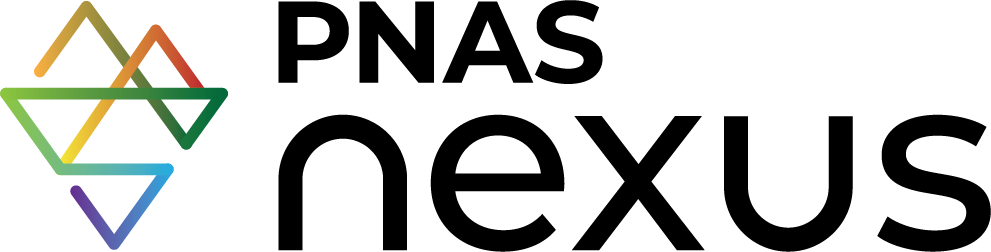

**Supplementary Material for**

**Mechanism of the Dual Action Self-Potentiating Antitubercular Drug Morphazinamide**

Lev Ostrer^1^†, Taylor A. Crooks^1^†, Michael D. Howe^1^, Sang Vo^1,2^, Ziyi Jia^1^, Pooja Hegde^2^, Nathan Schacht^1^, Courtney C. Aldrich^2^, Anthony D. Baughn^1^*

^1^Department of Microbiology and Immunology, University of Minnesota Medical School, Minneapolis, Minnesota USA

^2^Department of Medicinal Chemistry, College of Pharmacy, University of Minnesota Medical School, Minneapolis, Minnesota USA

†These authors contributed equally to this work.

*Correspondence can be made to Anthony D. Baughn.

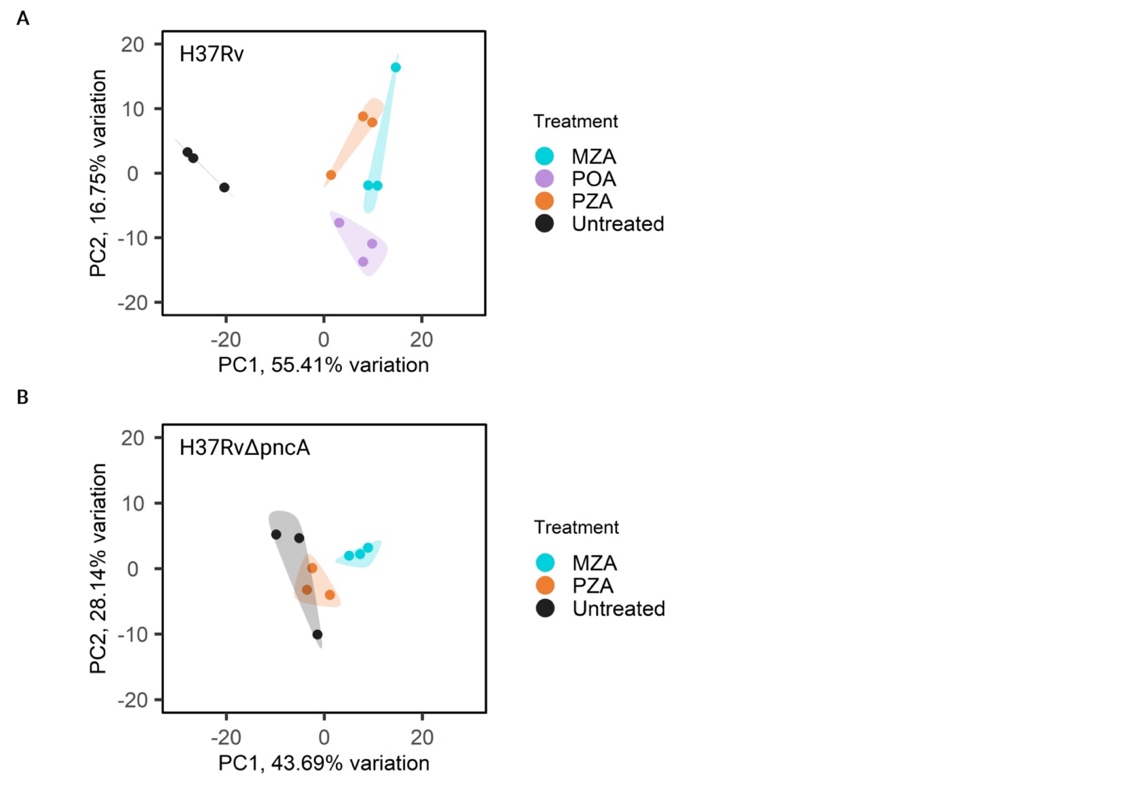

**Figure S1. PCA for RNA-seq treatment groups**. **A**, PCA plots from data from wildtype *M. tuberculosis* H37Rv cells treated with DMSO, PZA, POA or MZA. **B**, PCA plots from the *M. tuberculosis* H37Rv ∆*pncA* cells treated with DMSO, PZA or MZA. Each point represents an individual biological replicate
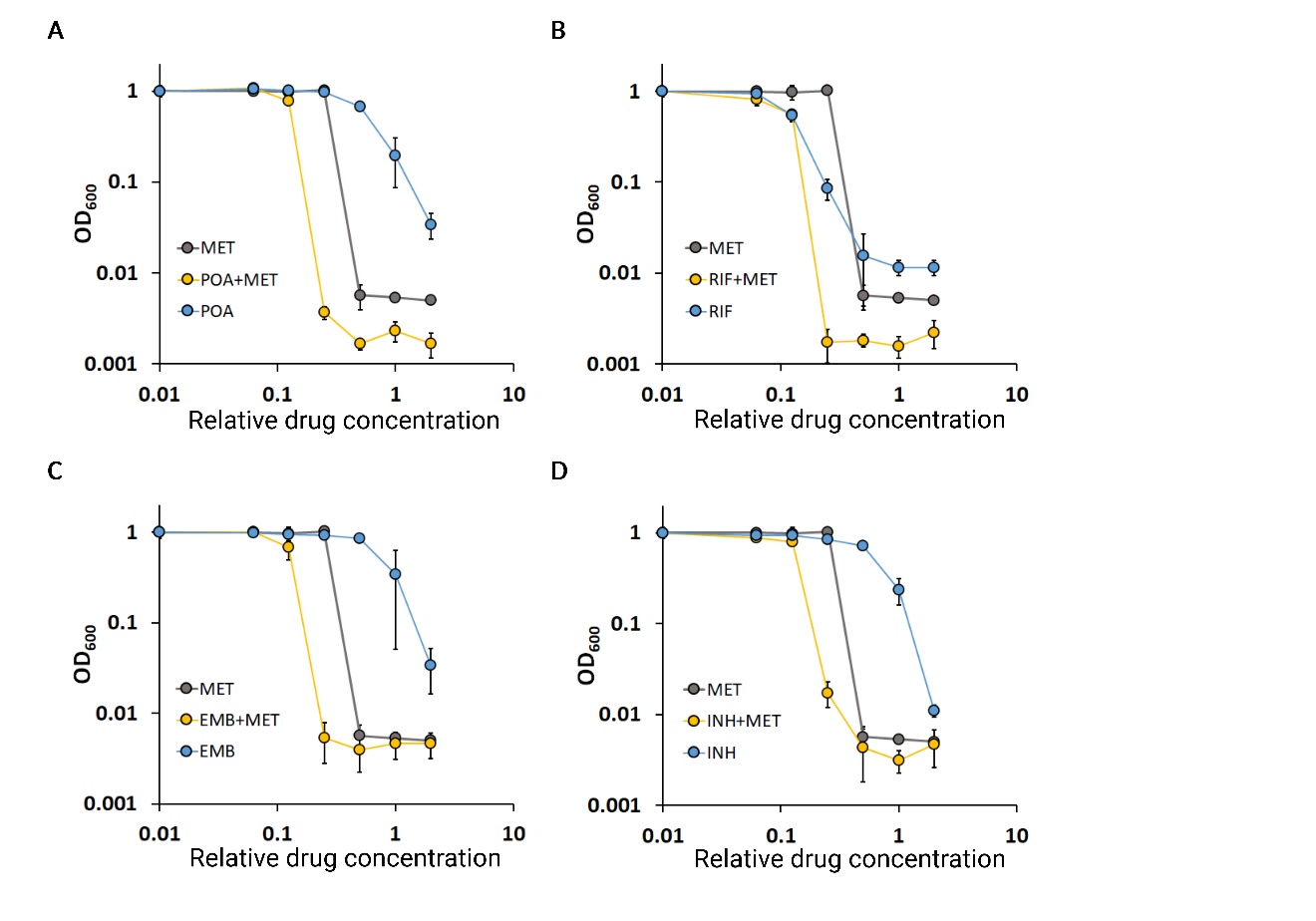

**Figure S2. DiaMOND Assay of Methenamine Interaction with First-Line TB Drugs**. H37Ra was used in each panel. Plots show exposure of *M. tuberculosis* H37Ra to a geometric series of methenamine (MET) alone and in combination with **A** POA**,** **B** Rifampicin (RIF), **C** Ethambutol (EMB), **D** Isoniazid (INH).

| **Table S1. PZA and MZA MIC values under potentiating and antagonizing conditions.** | | | |
| --- | --- | --- | --- |
|  |  | MIC in 7H9 media | |
| Strain | Condition | PZA (mM) | MZA (mM) |
| H37Ra | pH 5.8 | 0.13 | 0.07 |
| H37Ra | pH 5.8, pantothenate | >4.5 | 0.07 |
| H37Ra | pH 7 | >4.5 | 0.28 |
| H37Ra | pH 7, pantothenate | >4.5 | 0.56 |
| H37Rv | pH 5.8 | 0.25 | 0.07 |
| H37Rv | pH 7 | >4.5 | 0.28 |
| H37Rv Δ*pncA* | pH 5.8 | >4.5 | 0.28 |
| H37Rv Δ*sigE* | pH 5.8 | 2 | 0.14 |
| H37Rv Δ*sigE* | pH 7 | >4.5 | 0.28 |
| Experiments were performed using *M. tuberculosis* strains H37Ra and H37Rv and their derivatives. For potentiation of PZA action, pH of 5.8 was used. For antagonism of PZA action, pantothenate was used at a concentration of 0.23 mM. Strains H37Rv Δ*pncA* and Δ*sigE* were previously described (1) for their differing levels of PZA resistance. All assays were performed in biological triplicate with geometric mean displayed.  **References** | | | |

1.	J. M. Thiede *et al.*, Pyrazinamide Susceptibility Is Driven by Activation of the SigE-Dependent Cell Envelope Stress Response in *Mycobacterium tuberculosis*. *mBio* **13**, e0043921 (2021).
